# RIblast: An ultrafast RNA-RNA interaction prediction system for comprehensive lncRNA interaction analysis

**DOI:** 10.1101/077271

**Authors:** Tsukasa Fukunaga, Michiaki Hamada

## Abstract

Long non-coding RNAs (lncRNAs) play important roles in various biological processes. Although more than 58,000 human lncRNA genes have been discovered, most known lncRNAs are still poorly characterised. One approach to understanding the functions of lncRNAs is the detection of the interacting RNA target of each lncRNA. Because experimental detection of comprehensive lncRNA-RNA interactions are difficult, computational prediction of lncRNA-RNA interactions is an indispensable technique. However, the high computational costs of existing RNA-RNA interaction prediction tools prevents their application to large-scale lncRNA datasets. Here, we present “RIblast”, an ultrafast RNA-RNA interaction prediction method based on the seed-and-extension approach. RIblast discovers seed regions using suffix arrays and subsequently extends seed regions based on an RNA secondary structure energy model. Computational experiments indicate that RIblast achieves a level of prediction accuracy similar to those of existing programs, but at speeds over 64 times faster than existing programs.

---

Long non-coding RNAs (lncRNAs) play integral roles in diverse biological processes including histone modification [1], transcriptional regulation [2] and sub-nuclear structure formation [3]. The dysfunctions of many lncRNAs are associated with severe diseases such as coronary artery disease, diabetes, and various cancers [4, 5], and thus elucidating lncRNA functions is an important research area in molecular biology. Although large-scale transcriptome analysis has revealed that more than 58,000 lncRNA genes are encoded by the human genome [6], most of these lncRNAs are still poorly characterised [7].

Sequence similarity search and RNA secondary structure similarity search have achieved substantial success in characterising the function of protein-coding genes and short non-coding RNAs, respectively [8, 9]. However, these strategies are unsuitable for inferring the function of lncRNAs because lncRNAs frequently lack sequence and structure conservation [10, 11]. In contrast, the identification of interaction partners for each lncRNA should be a powerful approach to determining functions because lncRNAs function by being assembled with other proteins or RNAs into various complex molecular machinery [12].

Several lncRNAs have been experimentally confirmed to regulate biological processes through their interactions with target RNAs. For example, Abdel-mohsen *et al.* [13] determined that lncRNA 7SL reduces p53 protein translation levels by binding TP53 mRNA. Similarly, Carrieri *et al.* [14] found that lncRNA Uchl1-AS regulates the translation level of Uchl1 mRNA through an RNA-RNA interaction. Gong and Maquat [15] discovered that lncRNA 1/2-sbsRNAs inhibit the translation of the interaction target RNA through a Staufen1-mediated mRNA decay process. These examples show that the identification of lncRNA-RNA interactions is an important step in characterising lncRNA functions.

Several sequencing-based technologies have been developed as methods for the experimental discovery of RNA-RNA interactions. RIA-seq [16] and RAP-RNA [17] can identify target RNAs attached to an anchored RNA using *in vivo* cross-linking and antisense oligonucleotide probes. Although these methods are outstanding technologies to exhaustively detect interaction targets of a specific lncRNA, repeating these experiments across many lncRNAs is extremely labour intensive. In contrast, PARIS [18], SPLASH [19], LIGR-seq [20] and MARIO [21] can comprehensively identify RNA-RNA interactions *in vivo* based on proximity ligation. However, the majority of the detected interactions have been related to ribosomal RNAs or small RNAs, and the number of identified lncRNA-RNA interactions has been limited. In addition, because most of the lncRNAs show tissue-specific expression patterns [6, 10], these experiments on various tissues or cell lines are necessary but they require quite hard work and are therefore impracticable. Since the detection of genome-wide lncRNA-RNA interactions exclusively through experiments is difficult, computational prediction of lncRNA-RNA interactions is an indispensable technique.

Szcześniak and Makalowska [22] predicted entire lncRNA-RNA interactions across the human transcriptome using a fast sequence similarity search without consideration of RNA secondary structure. However, benchmarking results of RNA-RNA interaction predictions showed that omitting consideration of RNA secondary structure information decreases prediction accuracy [23]. To date, many RNA-RNA interaction prediction tools that consider RNA secondary structure have been proposed, e.g. IntaRNA [24], RNAplex [25, 26] and RactIP [27], and can detect small RNA (sRNA) interactions with high accuracy. However, as these programs were designed for detecting sRNA interactions, the computational costs are too high to predict lncRNA interactions comprehensively. To predict a comprehensive lncRNA interactome with consideration of RNA secondary structure, Terai *et al.* [28] first roughly screened interaction candidates based on only sequence complementarity and then exhaustively predicted lncRNA interactions using IntaRNA. Although their approach effectively narrowed down interaction candidates, it still required extensive computational resources to utilise IntaRNA. Therefore, a much faster RNA-RNA interaction prediction program that considers RNA secondary structure is required for further progress in comprehensive investigations of lncRNA function.

In the present study, we developed an ultrafast RNA-RNA interaction prediction algorithm for comprehensive lncRNA interaction analysis. While previous RNA-RNA interaction prediction tools employ a Smith-Waterman algorithm-like method, our algorithm is based on the seed-and-extension approach, which is widely adopted in sequence homology search tools including BLAST [8]. We implemented this high-speed algorithm as a program named RIblast, which detects seed regions using query and database suffix arrays, and subsequently extends both ends of seed regions based on an RNA secondary structure energy model. While the prediction accuracies of RIblast were comparable to those of existing programs, RIblast was more than 64 times faster than existing tools.

## Results

### Overview of the RIblast algorithm

RIblast enumerates potentially interacting segments between a query RNA *x* and a target RNA *y*. RIblat uses two energies as the evaluation criteria to determine whether two segments, (*x*_*s*_ and *y*_*s*_) in sequences *x* and *y*, intermolecularly interact: accessible energy and hybridization energy. *Accessible energy* is the energy required to prevent the segments from forming intramolecular base pairs and can be calculated by utilising a partition function algorithm [29, 30]. Briefly, a segment with high accessible energies tends to not form intermolecular base pairs because the segment forms intramolecular base pairs (Fig. 1A). *Hybridization energy* is the free energy derived from intermolecular base pairs between two segments and can be calculated as the sum of stacking energies and loop energies in the formed base-paired structure based on a nearest-neighbour energy model (Fig. 1B). When calculating hybridization energies, intra-molecular base pairs are not taken into consideration. Here, we defined the *interaction energy* between two segments *x*_*s*_ and *y*_*s*_ as the sum of the accessible energy of *x*_*s*_, accessible energy of *y*_*s*_ and hybridization energy between *x*_*s*_ and *y*_*s*_. RIblast outputs two segments with a particularly low interaction energy as a detected RNA-RNA interaction. Note that RNAup [31], IntaRNA [24] and RNAplex-a [26] also predict RNA-RNA interactions based on this combination of hybrid energy and accessible energy, and each showed high prediction accuracies in a previous benchmarking test [23].

**Figure 1:**
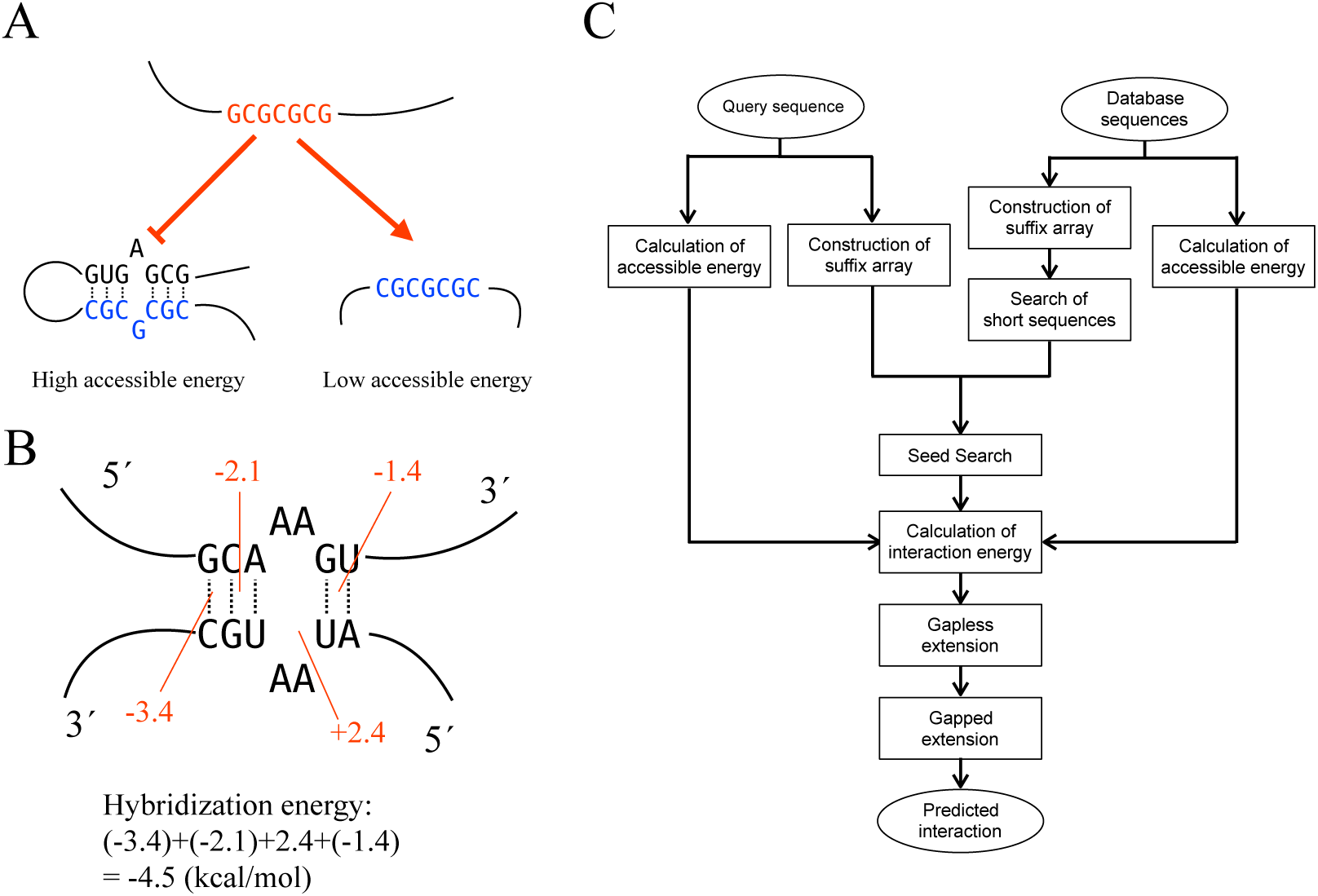
(A) A schematic illustration of the effect of accessible energies. While a segment with low accessible energy tends to form inter-molecular base pairs, a segment with high accessible energy tends not to form inter-molecular base pairs because such a segment tends to form intra-molecular base pairs. (B) Example of hybridization energy calculation. Hybridization energy can be calculated as the sum of stacking energies and loop energies in the formed base-paired structure. Generally, stacking energies stabilise RNA-RNA interactions but loop energies destabilize interactions. This calculation is based on Turner’s energy parameter. (C) Overview of the RIblast algorithm. The interaction energy is defined as the sum of hybridization energy and two accessible energies.

The accessible energy of each segment in an RNA sequence can be calculated with time complexity *O*(*NW*^2^) [32]. Here, *N* is the length of the input sequence and *W* is the constrained maximal distance between the bases that form base pairs. For all-to-all interaction predictions of lncRNAs, the calculation time of accessible energies scales linearly with the number of sequences. This is because accessible energies of an RNA sequence can be calculated independently of the other RNA sequences. On the other hand, the calculation of hybridization energy between two RNA segments is similar to the calculation of a local alignment score between two sequences [33]. Therefore, hybridization energy can be calculated based on a Smith-Waterman algorithm-like method with time complexity *O(NM)*, where *N* and *M* are the lengths of two input sequences; IntaRNA and RNAplex-a use this calculation approach. Unlike the calculation of accessible energies, the calculation of hybridization energies cannot be calculated from only an RNA sequence. Thus, the calculation time of hybridization energies is quadratic with the number of sequences when an all-to-all interaction prediction is conducted. This calculation is the obstacle to comprehensive lncRNA-RNA interaction prediction.

In the subject of local sequence alignment, the same problem was awaiting a solution, and a massive amount of research has been conducted to speed up the calculation of alignment scores. Seed-and-extension heuristic is one of the most successful approaches and has been adopted by many popular sequence alignment tools, such as BLAST [8], BLAT [34] and LAST [35]. This method first finds short matching regions, which are called seeds, between a query sequence and target sequence and subsequently extends alignments from both end points of the detected seeds. We recognised that the application of this approach to the calculation of hybridization energy should accelerate the computation speed considerably.

RIblast implements two major steps: database construction and an RNA interaction search. Fig. 1C shows the flowchart of the RIblast algorithm. In the database construction step, RIblast first calculates the accessible energy of each segment in the target RNA dataset using the Raccess algorithm [32]. To speed up calculation, RIblast calculates approximated accessible energies, as proposed in RNAplex-a [26], instead of exact accessible energies. Second, target RNA sequences are reversed and concatenated with delimiter symbols inserted between the two sequences. Third, a suffix array of the concatenated sequence is constructed. The suffix array is an efficient text-indexing data structure that comprises a table of the starting indices of all suffixes of the string in alphabetical order. It can be constructed in linear-time relative to sequence length [36, 37]. Fourth, in order to speed-up the RNA interaction search, search results of short strings are exhaustively pre-calculated. Then, the approximated accessible energies, concatenated sequences, suffix array and search results of short strings are stored in a database.

In the RNA interaction search step, RIblast first calculates approximated accessible energies and constructs a suffix array for a query RNA sequence. Second, RIblast finds seed regions whose hybridization energy is less than a threshold energy level *T*_1_ based on two suffix arrays of the query and the database. To efficiently enumerate seed regions, we used the modified algorithm of the seed search method of GHOSTX [38], which is a sequence homology search tool that is approximately 100 times faster than BLAST. Third, the interaction energies of the detected seed regions are calculated by summation of hybridization energy and two accessible energies. In this step, RIblast removes seed regions whose interaction energies exceed 0 kcal/mol. Fourth, RIblast extends interactions from seed regions without a gap. If RIblast extends the threshold length *Y* from the length requiring the minimum interaction energy in the extension but the minimum interaction energy has not been updated, then RIblast terminates the gapless extension. Fifth, the interactions that fully overlap with other interactions are removed. In addition, those interactions with interaction energies exceeding the threshold energy *T*_2_ are also excluded. Note that no interactions are removed if *T*_2_ is set to 0 kcal/mol, and lower *T*_2_ values cause faster computation speed with lower prediction accuracy. Finally, RIblast extends interactions from seed regions with a gap. As in the gapless extension step, if RIblast extends the threshold length *X* from the length requiring the minimum interaction energy in the extension but the minimum interaction energy has not been updated, then RIblast terminates the gapped extension. Further details of the algorithm are given in the Methods section. The source code of RIblast is freely available at https://github.com/fukunagatsu/RIblast.

### Evaluation of basepair prediction performance on bacterial sRNA dataset and fungal snoRNA dataset

We assessed the performance of RIblast using three evaluation methods. First, we investigated base pair prediction performance by evaluating whether programs predict correct base pairs between two RNAs with experimental interaction evidence. We used 109 validated bacterial sRNA-mRNA pairs and 52 validated fungal snoRNA-rRNA pairs as the evaluation dataset, which were constructed by Lai and Meyer [23] for the purpose of benchmarking RNA-RNA interaction predictions. To compare the performance of RIblast with other tools, we evaluated the base pair prediction performances of IntaRNA and RNAplex-a, which are the best performing current tools [23]. As the energy parameter characterising RNA secondary structures, we used two energy parameters, Turner’s energy parameter [39] and Andronescu’s BL_*_ energy parameter [40]. Because IntaRNA did not have an option to change the energy parameter, we used only the default Turner’s energy parameter in the IntaRNA evaluation. We used three accuracy measures: true positive rate (TPR), positive prediction value (PPV) and Matthews correlation coefficient (MCC). Positive base pairs were experimentally validated intermolecular base pairs [23]. The values of the adjustable parameter *T*_1_ and *X* in RIblast were determined based on the base pair prediction performance of the bacterial sRNA-mRNA dataset. The performances of various *T*_1_ and *X* values were investigated, and the parameter set that yielded the best performance was adopted (Supplementary Table S1 and S2). These determined values of *T*_1_ and *X* were used in the following analyses. *T*_2_ was set to 0 kcal/mol in this evaluation.

Tables 1 and 2 show the evaluation results of base pair prediction performance. For the bacterial sRNA-mRNA dataset, RIblast with Andronescu’s energy parameter achieved the best PPV (0.73) and MCC (0.67) performance. The best TPR score was obtained by IntaRNA (0.66). For the fungal snoRNA-mRNA dataset, RNAplex-a with Andronescu’s energy parameter was the best performing tool according to all three accuracy measures (TPR, 0.74; PPV, 0.69; MCC, 0.71), and was followed by RIblast using Andronescu’s energy parameter (TPR, 0.66; PPV, 0.60; MCC, 0.62). In both datasets, tools using Andronescu’s energy parameter showed superior performance to the same tool with Turner’s energy parameter.

**Table 1:** The results of base pair prediction performance on the bacterial sRNA dataset

| Program | TPR | PPV | MCC |
| --- | --- | --- | --- |
| IntaRNA | **0.66** | 0.61 | 0.62 |
| RNAplex-a (Turner) | 0.63 | 0.56 | 0.58 |
| RNAplex-a (Andronescu) | 0.60 | 0.68 | 0.63 |
| RIblast (Turner) | 0.58 | 0.66 | 0.61 |
| RIblast (Andronescu) | 0.63 | **0.73** | **0.67** |

The columns correspond to the three evaluation criteria: TPR, PPV and MCC. The rows indicate the performance of each program. The bold values are the highest scores in each column.

**Table 2:** The results of base pair prediction performance on the fungal snoRNA dataset

| Program | TPR | PPV | MCC |
| --- | --- | --- | --- |
| IntaRNA | 0.61 | 0.53 | 0.56 |
| RNAplex-a (Turner) | 0.56 | 0.49 | 0.52 |
| RNAplex-a (Andronescu) | **0.74** | **0.69** | **0.71** |
| RIblast (Turner) | 0.57 | 0.49 | 0.53 |
| RIblast (Andronescu) | 0.66 | 0.60 | 0.62 |

The columns correspond to the three evaluation criteria: TPR, PPV and MCC. The rows indicate the performance of each program. The bold values are the highest scores in each column.

### Evaluation of transcriptome-wide target prediction accuracy on bacterial sRNA dataset

Second, we evaluated bacterial sRNA target prediction performance by validating whether the predicted interaction energies of positive sRNA-mRNA interactions are lower than those of negative sRNA-mRNA interactions. This evaluation method was originally proposed by Richter and Backofen [41]. We used 64 experimentally validated interactions in *E. coli* as positive data. As negative data, we used all non-positive interactions in all-to-all interactions between 18 sRNAs and all 4319 *E. coli* mRNAs. We sorted mRNAs for each sRNA by minimum interaction energy. Then, we plotted ROC-like curves whose *x*- and *y*-axes were the number of true positive predictions and the total number of target predictions per sRNA, respectively. The parameter *T*_2_ was also set to 0 kcal/mol in this evaluation.

Fig. 2 shows the bacterial sRNA target prediction performance. The best performing tool was RNAplex-a with Andronescu’s energy parameter. The prediction performance of RIblast with Turner’s energy parameter was slightly lower, but RIblast with Andronescu’s energy parameter showed similar performance to the other programs.

**Figure 2:**
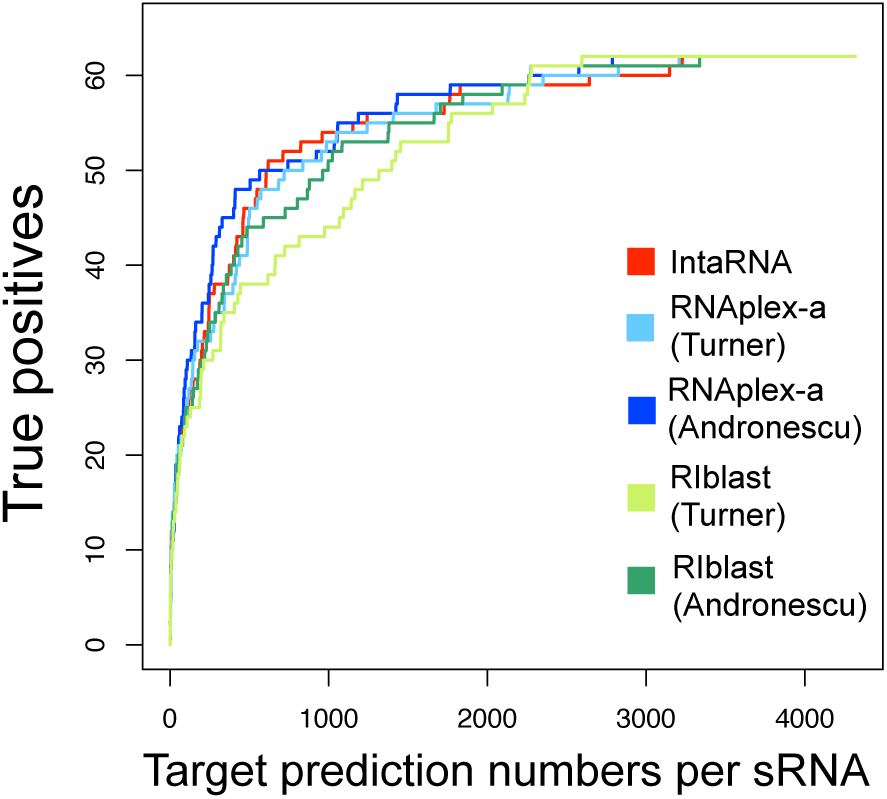
The performance of bacterial sRNA target prediction. The *x*- and *y*-axes represent target prediction numbers per sRNA and true positives, respectively. Red, sky blue, blue, light green and green colours represent the performances of IntaRNA, RNAplex-a (Turner), RNAplexa (Andronescu), RIblast (Turner) and RIblast (Andronescu), respectively. The best performing tool was RNAplex-a with Andronescu’s energy parameter. RIblast with Andronescu’s energy parameter exhibited performance similar to those of the other programs.

### Evaluation of human lncRNA TINCR target prediction accuracy

Third, we validated human lncRNA target prediction performance by comparing predicted interactions of human lncRNA TINCR with interactions experimentally validated by RIA-seq [16]. We used the same dataset and evaluation method as Terai *et al.* [28]. The dataset was composed of 5195 target RNAs (including both mRNAs and lncRNAs) and 1062 RNAs among them that interact with TINCR at one or more interacting segments. The target RNAs that have more interacting segments are more likely to be TINCR-interacting RNAs. As positive data, we used RNAs that at least had a threshold number of the interacting segments. When this threshold was set to 1, 2, 3, 4 and 5 interactions, the numbers of positive data were 1062, 434, 191, 104 and 65, respectively. Instead of comparing RIblast to IntaRNA or RNAplex-a, we compared the performance of RIblast with those of the pipeline by Terai *et al.* [28] and LAST [35], a fast local alignment tool. This is because lncRNA target predictions by IntaRNA and RNAplex-a have heavy computational costs. LAST was used by Szcześniak and Makalowska to make comprehensive human lncRNA-RNA interaction predictions [22]. We sorted target RNAs based on the minimum interaction energy among all predicted interactions in the target RNA (denoted by MINENERGY) or the sum of the interaction energies that are lower than some threshold value in the target RNA (denoted by SUMENERGY). Then, we calculated area under the receiver operating characteristic curve (AUROC) scores using the pROC R package [42].

Supplementary Table S3 shows AUROC results for MINENERGY sorting. LAST, the pipeline by Terai *et al.* [28] and RIblast exhibited performances that were similar to each other in this case. On the other hand, Fig. 3 and Supplementary Table S4-6 show AUROC scores for SUMENERGY sorting. This result illustrates that SUMENERGY sorting performs better than MINENERGY sorting among all methods. This result is consistent with at least one previous study [28]. In addition, for SUMENERGY sorting, RIblast achieved higher AUROC scores than the other methods for any threshold number of interacting segments. Unlike the evaluation of base pair prediction or sRNA target prediction performance, there was no difference in performance between Turner’s and Andronescu’s energy parameters. Finally, to obtain the appropriate parameter *T*_2_, we investigated the influence of *T*_2_ on TINCR target prediction accuracy (Supplementary Tables S7-8). These results show that the accuracy was robust to the *T*_2_ parameter setting. We set *T*_2_ to −6 and −4 when the energy models were Turner’s and Andronescu’s energy parameters, respectively.

**Figure 3:**
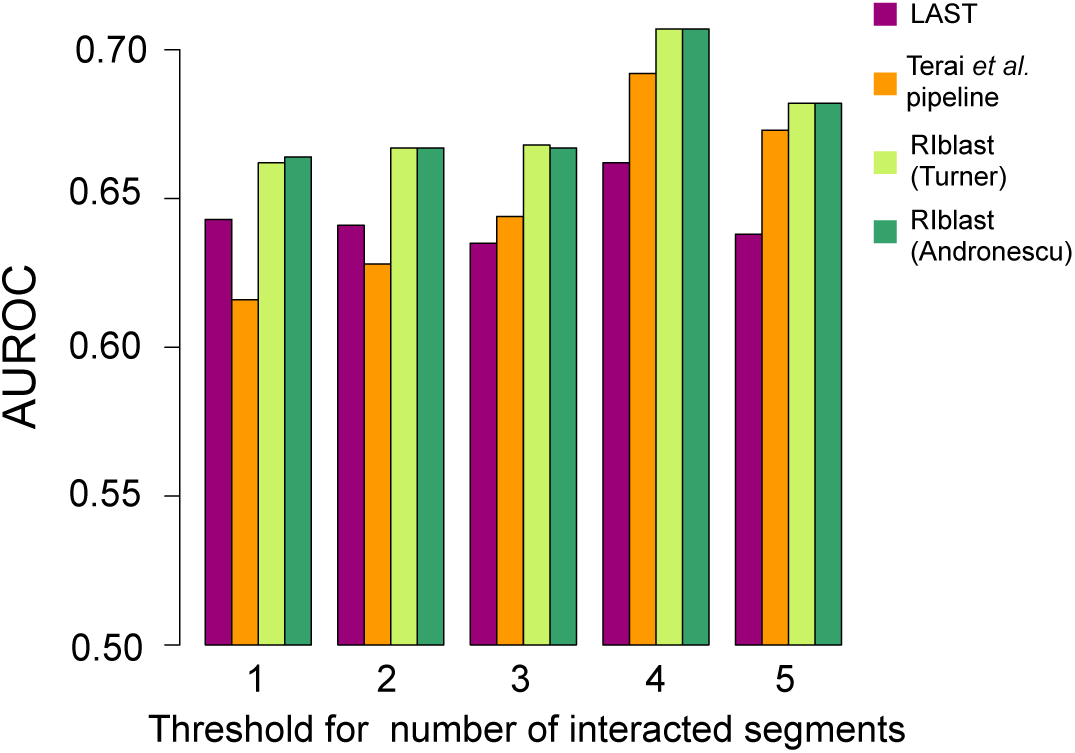
The performance of human lncRNA TINCR target prediction. The *x*-axis represents the threshold number of interacting segments in the positive data. The *y*-axis represents the area under the receiver operating characteristic curve (AUROC) score. Purple, orange, light green and green colours represent the performances of LAST, the Terai *et al*. pipeline, RIblast (Turner), and RIblast (Andronescu), respectively.

### Evaluation of running time

We finally evaluated the computational speed of RIblast by comparing its run time with the times required for IntaRNA, RNAplex-a, and the pipeline by Terai *et al.* [28]. We excluded the joint secondary structure prediction step using RactIP [27] in the Terai *et al.* [23] pipeline because this step does not affect interaction prediction accuracy. The calculation time for RNAplex-a included the run time of accessibility calculation by RNAplfold [43], and that of RIblast includes both the execution time of the database construction step and the RNA interaction search step. The query and target sequences were randomly selected from human lncRNAs and mRNAs in Gencode version 24, respectively [44]. Then, all-to-all interaction predictions between query and target sequences were conducted. The computation was performed on an Intel(R) Xeon E5 2670 2.6GHz CPU with 4 GB of memory. Table 3 shows the computational times depended on the dataset size for each software tool. In all cases, RIblast was much faster than the other programs. As the dataset size increased, the speed advantage over the other programs became quite large. In particular, when the dataset consisted of 500 lnRNAs and mRNAs, RIblast was 64-fold and 73-fold and faster than the Terai *et al.* pipeline and RNAplex-a, respectively (Table 4).

**Table 3:** The results of the run time evaluation on partial human lncRNA and mRNA datasets

| Program | The number of lncRNAs and mRNAs |  |  |  |  |
| --- | --- | --- | --- | --- | --- |
|  | 5 | 10 | 50 | 100 | 500 |
| IntaRNA | 59m 04s | 3h 30m 17s | - | - | - |
| RNAplex-a | 2m 34s | 10m 37s | 4h 20m 42s | 17h 56m | 19d 20h 53m |
| Terai *et al.* pipeline | 1m 02s | 3m 08s | 2h 26m 43s | 14h 50m | 17d 09h 44m |
| RIblast | 28s | 50s | 5m 33s | 16m59s | 6h 26m |

The columns correspond to the number of lncRNAs assessed in the dataset. The rows indicate the run times of each program. The symbol “-” indicates that we did not investigate the computational speed for a particular combination of dataset size and program because the calculation time was prohibitively long.

**Table 4:** Calculation speed ratios of RIblast to those of the other programs on partial human lncRNA and mRNA datasets

| Program | The number of lncRNAs and mRNAs |  |  |  |  |
| --- | --- | --- | --- | --- | --- |
|  | 5 | 10 | 50 | 100 | 500 |
| IntaRNA | 126.6 | 252.3 | - | - | - |
| RNAplex-a | 5.5 | 12.8 | 47.0 | 63.4 | 74.117 |
| Terai *et al.* pipeline | 2.2 | 3.8 | 26.4 | 52.4 | 64.9 |

The columns correspond to the number of lncRNAs in the dataset. The rows indicate the run time ratio of each program to RIblast. The symbol “-” indicates that we did not investigate the computational speed for a particular combination of dataset size and program because the calculation time was prohibitively long.

## Discussion

In this study, we developed a novel RNA-RNA interaction prediction algorithm based on the seed-and-extension approach and implemented it as RIblast. RIblast showed comparable accuracies to the current tools with the best base pair prediction performance and sRNA target prediction performance, and RIblast also showed superior performance to existing tools in human lncRNA TINCR target prediction. Moreover, RIblast is much computationally faster than the other programs assessed. These results strongly suggest that the seed-and-extension approach is effective for accelerating RNA-RNA interaction predictions, and RIblast is the top choice as a tool for comprehensive lncRNA-mRNA interaction prediction.

We used an interaction energy cutoff to exclude likely incorrect predictions in this research, but this method may be highly arbitrary. As such, we should ultimately determine the reliability of the predicted interactions based on a statistical score like the e-values generated by BLAST. Rehmsmeier *et al.* [45] developed a calculation method for the statistical significance of predicted RNA-RNA interactions. However, their calculation method cannot be applied to our software directly because their interaction prediction method did not consider the effect of accessible energies. Therefore, we need to develop a novel e-value calculation method for RIblast’s predicted interactions.

Although Hajiaghayi *et al.* reported that the accuracy of RNA secondary structure prediction with Andronescu’s energy parameter outperforms those that use other energy parameters [46], our research provides the first report that Andronescu’s energy parameter also delivers superior performances compared with Turner’s energy parameter in small RNA-RNA interaction predictions. Currently, major miRNA target prediction tools, such as miRanda [47] and TargetScan [48], and snoRNA target prediction tools, such as RNAsnoop [49], utilise Turner’s energy parameter. The application of Andronescu’s energy parameter to these programs may easily improve their target prediction accuracy.

RIblast efficiently calculates RNA-RNA interaction predictions, but further acceleration is an essential task because the number of lncRNA is increasing daily. Considering that the seed-and-extension approach greatly contributes to the acceleration of RNA-RNA interaction predictions, other acceleration techniques in sequence homology search may be effective for the acceleration of RNA-RNA interaction predictions. Specifically, algorithm parallelization is a promising technique. At present, many parallelization methods based on GPGPU [50, 51, 52], MPI [53] and SIMD [54] have been proposed for sequence homology search and have successfully speed up calculation.

While typical mRNAs tend to be localised in the cytoplasm, typical lncRNAs tend to be localised in the nucleus [55]. This tendency may suggest that lncRNAs exert their gene regulatory functions by interacting with nascent pre-mRNAs [17]. Thus, comprehensive interaction prediction between lncRNAs and pre-mRNAs is a fascinating research topic, but the current version of RIblast cannot be applied to this task. This is because the accessible energy calculation of pre-mRNAs by the Raccess algorithm is computationally difficult for long RNA sequences. For this purpose, we will integrate the ParasoR algorithm [56], which can calculate accessible energies for quite long RNAs on a computer cluster, with RIblast.

The evolution of lncRNA is a hot topic in RNA biology [57]. Although the majority of lncRNAs are lineage-specific, a thousand human lncRNAs have homologs with conserved short sequence regions [11]. In addition, Ngueyn *et al.* revealed that experimentally validated RNA-RNA interaction sites are evolutionarily conserved [21]. These results suggest that the interaction relationships between lncRNA and RNA are widely conserved among species. We aim to validate this hypothesis by comparing RIblast-based lncRNA interactome networks between species.

## Methods

### The method for calculating accessible energy

We presumed that the conformation distribution of RNA secondary structures of an RNA sequence is represented by the Boltzmann distribution. We defined *x*[*i*.. *j*] as a segment from position *i* to position *j* in an RNA sequence *x*. Here, the accessible energy *E*_*acc*_(*i*, *j*) that is required to make the segment form a single-stranded structure is given by

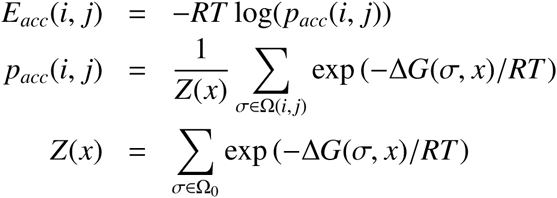

where ∆*G*(*σ, x*) represents the Gibbs free energies of the given structure *σ* on the sequence *x*, *R* represents the gas constant and *T* represents the absolute temperature (we used *T* = 310.15 K in this study). Ω_0_ represents the set of all possible secondary structures of *x*, and Ω(*i*, *j*) is the set of all possible secondary structures that the segment *x*[*i*.. *j*] forms in single-stranded structure. Hence, *p*_*acc*_(*i*, *j*) is the probability that the segment *x*[*i*.. *j*] is single-stranded. For a fixed segment length, Raccess can calculate accessible energies of all segments with *O*(*NW*^2^) using dynamic programming, where *N* is the sequence length and *W* is the constraint of maximal distance between the bases that may form base pairs. In this research, we uniformly set *W* to 70 as a research benchmark for RNA-RNA interaction prediction tools [23].

However, RIblast requires the accessible energies of segments with arbitrary length, and the exhaustive calculation is computationally expensive. Therefore, RIblast uses approximated accessible energies 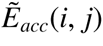 instead of *E*_*acc*_(*i*, *j*). This method was proposed in RNAplex-a [26]. 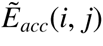 was defined as follows:

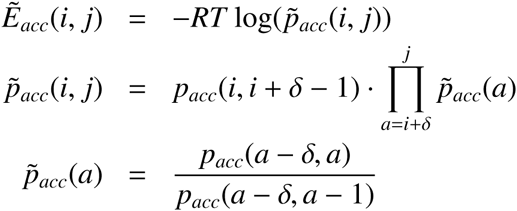

By this approximation, we only have to calculate the accessible energies of segments with length *δ* and *δ* + 1. In addition, by restricting the minimum length of seeds to *δ*, we need not calculate accessible energies of segments whose length is less than *δ*. Note that when the segment length is *δ* or *δ* + 1, the approximate accessible energy 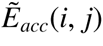 becomes the exact accessible energy *E*_*acc*_(*i*, *j*). In this research, *δ* was set to 5.

### Seed search

The seed design strongly influences the accuracy and calculation speed of the program. BLAST searches seeds with a fixed length, but this method is unsuitable for RNA-RNA interaction search. For example, in Andronescu’s energy model, the hybridization energy of a 6-mer seed consisting of only G-C base pairs is about −10 kcal/mol, but that consisting of only G-U base pairs is about −1 kcal/mol. The large difference in hybridization energies between seeds of the same length should depress the performance of tools. Therefore, RIblast adopts score-based seeds, as proposed in GHOSTX [38]. Our score-based seeds were defined as the perfect base-pairing region whose hybridization energy is less than the threshold energy *T*_1_ and length is at least *δ*. Note that the seed search step takes hybridization energy into consideration, but does not consider the accessible energies of the two segments.

RIblast detects seeds using a depth-first search. Supplementary Figure S1 shows the schematic illustration of the seed search. First, RIblast searches for a single inter-molecular base pair such as G-C. If this pair is found in the query and the database, then RIblast extends the base pair by one base pair as GG-CC, GC-CG,…, GU-CG and then checks whether these extended strings are found in the query and the database. If the extended strings are detected and meet the conditions for score-based seeds, then RIblast stores the string pair as a seed. If extended strings are detected but do not meet the conditions for score-based seeds, then RIblast extends the strings by one base pair again and repeats this step. If extended strings are not detected, then the extension is stopped. To avoid overly long seeds, we restricted the max seed length *length*_*max*_ (we set this parameter to 20 in this study). Supplementary Figure S2 shows the pseudo-code of the RIblast seed search algorithm. Here, *S*_*q*_ and *S*_*db*_ represents the query RNA sequence and the reversed and concatenated database RNA sequence, respectively. *S A*_*q*_ and *S A*_*db*_ are the suffix arrays of *S*_*q*_ and *S*_*db*_, respectively. *seed*_*q*_ and *seed*_*db*_ represent the temporary seeds for the query and database, respectively. *sp*_*q*_, *ep*_*q*_, *sp*_*db*_ and *ep*_*db*_ are the indices of *S A*_*q*_ and *S A*_*db*_. The *S AS earchNextS tring* function returns the indices of the new extended string in a suffix array. If the string exists in the query and the database, then the returned *sp*(*sp*^′^) is smaller than the returned *ep*(*ep*^′^).

In order to accelerate this seed search step, we pre-calculate the indices of the strings whose length is shorter than *l* for a database of RNA sequence in the database construction step. The results of the short sequence search are used on the database sequences. Therefore, this binary search of the suffix array is needed only for the search of query sequences or long strings in the database sequence. In this research, we set *l* to 8.

### Extension

After the seeds are found in the query and the database, RIblast tries to extend interactions from both end points of these seed regions. The gapless extension is first conducted, and then the gapped extension is performed in a similar way to BLAST, LAST and GHOSTX.

RIblast first extends interactions without a gap from seed regions. If extended interactions have lower interaction energies than the present minimum interaction energy in this extension step, then RIblast updates the minimum interaction energy. Otherwise, extensions are repeated. If RIblast extends *Y* nucleotides from the length that requires the minimum interaction energy in this extension but the minimum interaction energy has not been updated, then RIblast terminates the gapless extension. In this step, we assume that the possible complementary bases always interacts with each other. After the gapless extension step, if two interactions {*S*_*q*_[*i*, *j*], *S*_*db*_[*k*, *l*]} and {*S*_*q*_[*i*^′^, *j*^′^], *S*_*db*_[*k*^′^, *l*^′^]} satisfy the conditions *i* ≤ *i*^′^, *j* ≥ *j*^′^, *k* ≤ *k*^′^ and *l* ≥ *l*^′^, then we exclude the later interaction. In addition, if the interaction energy of an interaction exceeds threshold *T*_2_, then we also remove the interaction. In this research, we set 5 to Y.

Next, RIblast tries to extend interactions with a gap. Like the gapless extension step, if the interaction energy of extended interactions is lower than the present minimum interaction energy in this extension step, then the minimum interaction energy is updated. If RIblast extends *X* nucleotides from the length that requires the minimum interaction energy in this extension but the minimum interaction energy has not been updated, then RIblast terminates the gapless extension. The calculation of the interaction energy of extended interactions is as follows (Supplementary Figure S3 shows the schematic illustration). Here, we regard {*S*_*q*_[*i*, *j*], *S*_*db*_[*k*, *l*]} as an interaction after gapless extension. In the extension towards the 5^′^ end of the query sequence (and 3^′^ end of the database sequence), RIblast calculates *E*_*int*_(*a*, *b*), which is the minimum interaction energy for sequences *S*_*q*_[*a*, *j*] and *S*_*db*_[*b*, *l*], as the following equation.

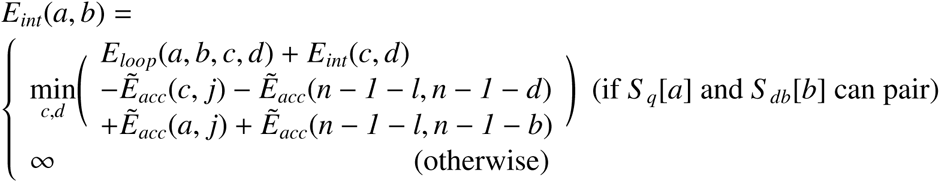

where *E*_*loop*_(*a*, *b*, *c*, *d*) indicates the free energy of the loop consisting of base pairs (*a*, *b*) and (*c*, *d*) and *n* is the sequence length of the database sequence. Here, *a* < *c* ≤ *i* < *j* and *b* < *d* ≤ *k* < *l* are satisfied. In addition, the internal loop size *c* − *a* + *d* − *b* is restricted to within *X*. Note that many RNA secondary structure prediction tools such as RNAfold [58] adopt this restriction of internal loop size. The extension in the opposite direction is calculated in the same manner. Dangling energies are added only after gapped extensions are finished.

### Repeat masking

Three types of repeat masking were implemented in RIblast: no-masking, soft-masking and hard-masking. The no-masking procedure treats repeats just like non-repeat sequence. The soft-masking procedure excludes repeat sequences in the seed search step, but considers them in the extension step. The hard-masking procedure completely ignores repeats. We used the soft-masking procedure to evaluate TINCR target prediction in order to match our study with previous research by Terai *et al.* [28]. For the other evaluations, as we did not use repeat masking tool, the type of repeat masking used did not affect the results.

### Method for evaluating base pair prediction performance

To evaluate the base pair prediction performance, we used 109 validated bacterial sRNA-mRNA pairs and 52 validated fungal snoRNA-rRNA pairs as datasets. The bacterial sRNA-mRNA interaction dataset was composed of 64 *E. coli* and 45 *Salmonella enterica* interactions as well as 18 query sRNAs and 82 target mRNAs. Following the benchmark research of Lai and Meyer [23], we used the sequences between 150 bp upstream and 150 bp downstream of each start codon as the target sequences. All fungal snoRNA-rRNA interactions in the dataset were *S. cerevisiae* C/D box interactions, and these interactions were between 43 snoRNAs and 2 rRNAs. For target rRNAs, full rRNA sequences (1800 nucleotide 18S rRNA and 3396 nucleotide 25S rRNA) were used. We compared the performance of RIblast with those of IntaRNA and RNAplex-a. The command line options used for IntaRNA and RNAplex-a in the present study were the same as those used by Lai and Meyer in their benchmark research [23]. TPR, PPV and MCC were calculated for each RNA-RNA interaction, and the averaged scores were evaluated. The definitions of these three scores are as follows:

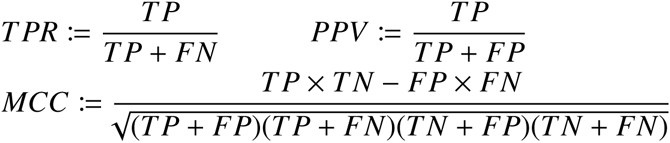

We discarded suboptimal predictions and only evaluated the minimum energy interactions. To determine the values of parameter *T*_1_ and *X*, we investigated the accuracy of 63 parameter combinations for each energy parameter. *T*_1_ is a threshold energy for score-based seed detection, and *X* is a threshold length for extension termination. The parameter combinations consisted of 7 *X* parameters and 9 *T*_1_ parameters. We adopted the parameter combination that yielded the highest MCC score. If there were several parameter combinations with the best performance, we adopted the smallest *X* and largest *T*_1_ parameter combination in order to accelerate computation. As a result, we set *X* and *T*_1_ to 18 and −10.0, respectively, when we using Turner’s model, and we set *X* and *T*_1_ to 16 and −6.0, respectively, when we used Andronescu’s model.

### Method for evaluating sRNA target prediction performance

We evaluated the sRNA target prediction performance by predicting all-to-all interactions between 18 sRNAs and all 4319 *E. coli* mRNAs. As target mRNA sequences, we used sequences between 150 base-pairs (bp) upstream and 50 bp downstream from each start codon. This sequence length setting is the same as that used by Terai *et al.* [28]. The sequence data were downloaded from NCBI (http://www.ncbi.nlm.nih.gov/nuccore/NC_000913). We used 64 experimentally validated interactions as positive data, which were also used to evaluate base pair prediction performance. Only the predicted interaction with the minimum interaction energy was evaluated.

### Evaluation method for human lncRNA TINCR target prediction accuracy

To evaluate the TINCR target prediction performance, we used an RIA-seq-based TINCR interaction dataset [16]. The simple repeat regions were masked by TANTAN [59] with the default options. We compared the performance of RIblast with that of LAST [35] and the pipeline by Terai *et al.* [28]. In LAST, we set G-C, A-U and G-U match scores to 4, 2 and 1, respectively. The mismatch score, gap opening penalty and gap extension penalty were set to −6, −20 and −8, respectively. These parameter settings are the same as those used by Szcześniak and Makalowska [22]. We regarded the score of the detected alignment × (−1) as the interaction energy between the regions. The short summary of the Terai *et al.* pipeline is as follows. First, accessible energies were calculated by Raccess, and inaccessible regions were removed from the analysis. Second, pairs of complementary gapless subsequences were detected as interaction regions by LAST. Finally, the interaction energies of the interaction regions were calculated by IntaRNA.

To determine values for the parameter *T*_2_, we examined the dependence of accuracy decreases from AUROC scores of SUMENERGY on *T*_2_. We used AUC scores of −16 kcal/mol and −8.5 kcal/mol as interaction energy thresholds for SUMENERGY when the energy parameters were Turner and Andronescu parameters, respectively.

## Acknowledgments

This research was supported by the Japan Society for the Promotion of Science [grant numbers JP16J00129 and JP16H05879]. We thank Dr. Junichi Iwakiri for the helpful discussion, and Dr. Kun Qu and Dr. Paul A. Khavari for providing the TINCR RIA-seq dataset.

### Author Contributions

TF and MH designed the project. TF developed the algorithm and performed the analyses. TF and MH wrote the paper. Both the authors read and approved the final manuscript.

### Competing interests

The authors declare no conflict of interest.

